# Enhancing Knowledge Discovery from Cancer Genomics Data with Galaxy

**DOI:** 10.1101/089631

**Authors:** Marco A. Albuquerque, Bruno M. Grande, Elie J. Ritch, Selin Jessa, Martin Krzywinski, Jasleen K. Grewal, Sohrab P. Shah, Paul C. Boutros, Ryan D. Morin

## Abstract

We present a collection of Galaxy tools representing many popular algorithms for detecting somatic genetic alterations from cancer genome and exome data. We implemented methods for parallelization of these tools within Galaxy to accelerate runtime and have demonstrated their usability on cloud-based infrastructure and commodity hardware. Some tools represents extensions or refinement of existing toolkits to yield visualizations suited to cohort-wide cancer genomic analysis. For example, we present Oncocircos and Oncoprintplus, which generate data-rich summaries of exome-derived somatic mutation. Workflows that integrate several of these to perform some standard data integration and visualization tasks are demonstrated on a cohort of 96 diffuse large B-cell lymphomas, enabling the discovery of multiple candidate lymphoma-related genes that have not been reported previously.

## Background

An inherent problem in the application of genomics to understand the molecular aetiology of cancer is the multi-disciplinary skillset required for researchers to draw meaningful inferences from high-throughput biological data. With the rise in popularity of high-throughput DNA sequencing, the bottleneck for novel discovery has shifted from data generation to data analysis and interpretation. Although myriad algorithms have been developed to efficiently analyze increasingly large datasets, these are often tailored for technically-inclined users. Specifically, they are typically run at the command line; their operation requires the use of cryptic parameters; and their installation is often burdensome. Data flow between tools is also often non-trivial and, owing to a paucity of data standards, can involve error-prone data manipulation and re-formatting steps. Combined with a necessity for high-performance computational hardware to run many such tools efficiently, this produces a tremendous barrier for new users.

There exists a handful of options that address this predicament in genomics as a whole. Tools that automate pipeline development such as Kronos ^1^, Nextflow ^2^ and Snakemake ^3^ can satisfy the needs of more technically savvy users. Alternatively, graphical user interfaces (GUIs)—which are generally lacking in the field of bioinformatics—aid in users learning the utility of the associated with command-line interfaces but typically do not scale to large data sets. Examples of genomics tools offering web-accessible GUIs include `BLAST` ^4^, `VAGUE` ^5^ and `limmaGUI` ^6^. However, beyond an inability to scale, web-based utilities pose several issues, including design inconsistency, redundant efforts in interface development and the inability to automatically link individual tasks into pipelines or workflows. Ideally, any reduction in the barriers associated with running individual algorithms passing data between software tools should accelerate analytical tasks and reduce the risk of errors.

To overcome this, GUI-enabled software for automating pipeline development has been created to improve the reproducibility, accessibility and transparency of running genomic analyses ^7^. Examples of these include Galaxy ^8^, Taverna ^9^, Pegasus ^10^ and commercial software packages such as Geneious ^11^. In particular, the Galaxy project offers many attractive features for this goal while remaining open-source. Namely, Galaxy boasts extensive documentation; support for automatic tool installation; the ability to instantiate public or private “cloud clusters” by leveraging CloudMan ^12^; and is as a whole supported by a vibrant community that provides ongoing development to the software. Although algorithms for handling high-throughput sequence data are increasingly being added to Galaxy, there currently remains a general lack of tools and workflows specifically tailored to perform common tasks involved in analyzing cancer genome and exome sequence data. Here, we have begun to address this issue by adapting many of the popular tools for analyzing cancer genome and exome data for Galaxy and made these publicly available (https://github.com/morinlab/tools-morinlab). We demonstrate the utility of some of these by applying custom workflows to a large cohort of diffuse large B-cell lymphoma patients (n=96) and uncover new candidate lymphoma-related genes.

## Results

### Implementing cancer genomics tools in Galaxy

We produced a comprehensive toolkit comprising a suite of complementary tools and workflows that perform many of the routine analytical tasks in cancer genomics. These include several methods for detecting (“calling”) somatic single nucleotide variants (SNVs), copy number variations (CNVs) and structural variations (SVs) in tumour-normal pairs. Additional tools were developed to perform the many auxiliary steps helper functions that allow these tools to be linked and applied generically, such as bam and text file pre‐ and post-processing, manipulating and converting file formats; variant annotation; identification of significantly mutated genes; and multiple visualization utilities for performing exploratory analysis and cohort-level data summarization. The tools and helper functions are detailed in Table 1 and Supplementary Table S1, respectively. Details on the implementation of tools is included in the Methods section and can be further obtained from the latest versions hosted in our repository (https://github.com/morinlab/tools-morinlab).

**Table 1:** Main tools currently comprising the cancer genomics toolkit. Tools representing existing or extended analysis approaches are shown above. For a current list of tools available, refer to the github repository: https://github.com/morinlab/tools-morinlab

| Tool | Category | Reference |
| --- | --- | --- |
| mutationSeq | SNV detection | 29 |
| Strelka | SNV and indel detection | 30 |
| SomaticSniper | SNV detection | 31 |
| RADIA | SNV detection | 32 |
| VarDict (Java) | SNV detection | 33 |
| DELLY | SV detection | 34 |
| LUMPY | SV detection | 35 |
| Pindel | SV and indel detection | 36 |
| Manta | SV detection | 37 |
| Sequenza | CNV detection | 38 |
| TITAN | CNV detection | 39 |
| Ensembl VEP | SNV Annotation | 40 |
| PyClone | Clonal structure | 41 |
| EXPANDS | Clonal structure | 42 |
| MutSigCV | Significantly Mutated Genes | 43 |
| Oncodrive-FM | Significantly Mutated Genes | 44 |
| GISTIC | Significantly Mutated Genes | 45 |
| Maftools (oncostrip, oncodriveclust, oncoplot, trinucleotide plot, MAF summary, rainfall plot, lollipop plot) | Visualization, significantly mutated genes, mutation signatures | 15 |
| Oncocircos | Visualization | 20 |
| Oncoprintplus | Visualization | 46 |

There are numerous algorithms available to perform standard analytical tasks such as variant calling and CNV detection, each offering different balances of usability, computational efficiency and accuracy. As such, selection of ideal tools and parameters is non-trivial. We implemented tools representing some of the more commonly cited options and include many that performed favorably in ICGC-TCGA DREAM challenges ^13^. As each tool can be configured with a number of parameters, which can be tuned for improved accuracy, we leverage results from the DREAM challenge to assist in selecting the more accurate algorithms and in setting sensible default parameters ^14^. We have also released numerous workflows that run some of the more complicated pieces of software that relies on many dependencies and that perform some routine analytical and visualization tasks as detailed and illustrated with the real-world worked examples below. Example workflows that demonstrate our new approach to perform parallelization in Galaxy are also included (Figure 1). As these migrate to the public Galaxy tool shed, we encourage ongoing testing and parameter optimization and community-driven refinement and expansion of this toolkit such that it may eventually constitute a hub for boilerplate cancer genome analyses.

**Figure 1.**
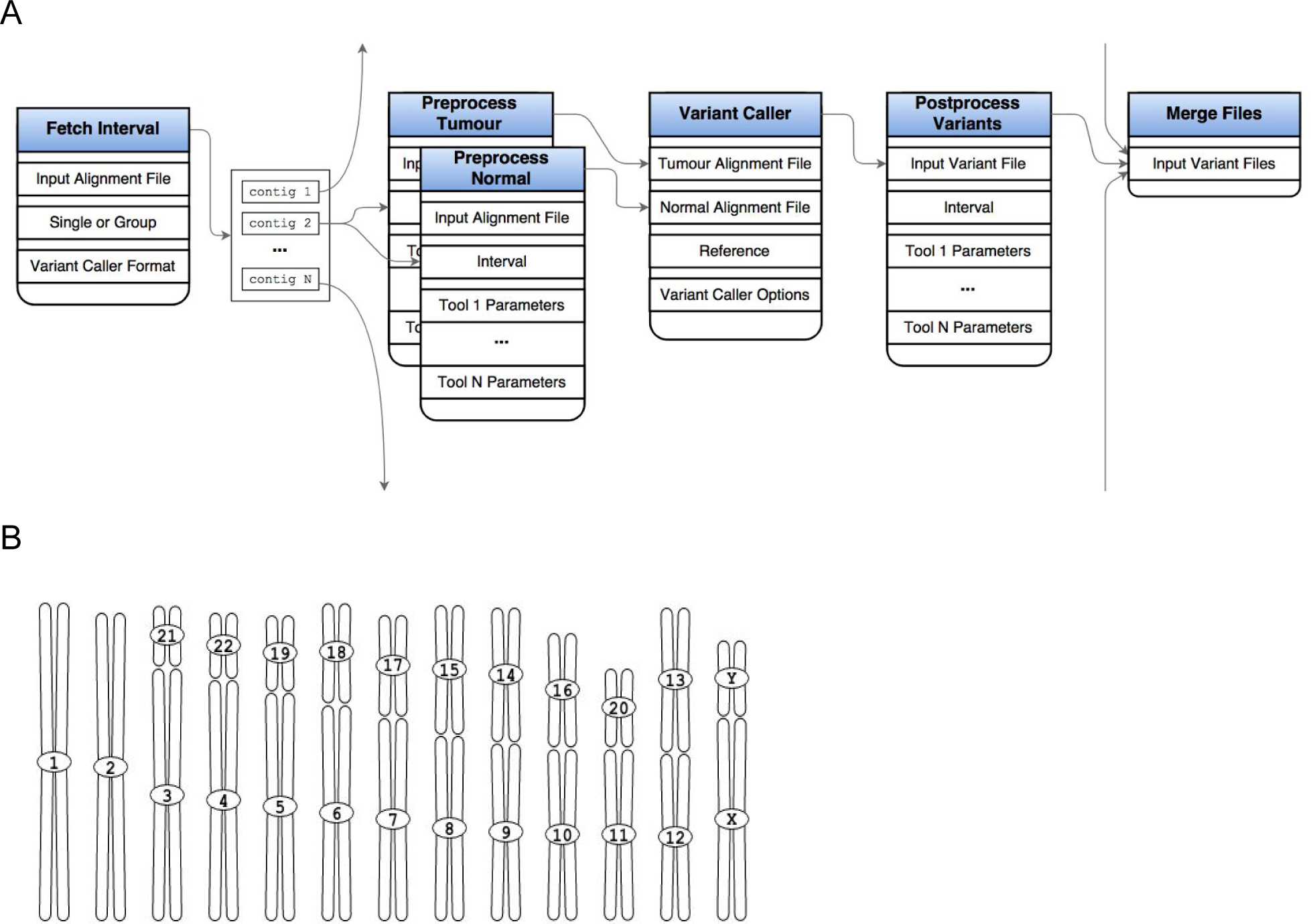
Parallelization in variant calling and other CPU-intensive processes. (A) An alignment file flows through to `fetch_interval`, which obtains all contigs in an alignment file. If parallelization is requested, multiple interval files are generated for each interval, otherwise a single file is created. Each dataset in the collection is treated as separate input to two instance of `preprocess`, which filters reads from the sequence alignment file for normal and tumour alignment files. These then pass to the variant caller. A `postprocess` tool filters and annotate variant calls based on tool-specific parameters and all final variants are merged and sorted in a single variant file. (B) We perform automatic interval selection to roughly balance the load on each variant-calling task. The algorithm combines regions (e.g. chromosomes) if their total length is less than the largest. In cloud-based settings, this reduces overhead associated with creating multiple unnecessary parallelized jobs as well as reducing the number of short-lived automatically added nodes. Importantly, we chose not to implement a sub-chromosomal interval selection algorithm to maintain intrachromosomal dependence required by some of the variant calling algorithms. Such an extension could be implemented for tools that lack this restriction.

### Identifying novel candidate lymphoma-related genes from exome data

An ultimate goal in cancer genome/exome analysis involves the identification of loci recurrently affected by copy number gain or loss and genes recurrently targeted by somatic mutations. There exist myriad tools to detect somatic SNVs and a growing number of options to derive high-quality copy number estimates from genome and, in less cases, exome data. We implemented workflows that perform the required annotation and pre-processing of raw mutation and copy number outputs from tools such as `Strelka` and `Sequenza`, respectively. We ran these two workflows on 96 tumor/normal pairs representing diffuse large B-cell lymphoma (DLBCL) patients. The SNV and indel calls were annotated and converted into the standard mutation annotation format (MAF) using `vcf2maf`. We subsequently analyzed the pooled mutation calls from the meta-cohort for recurrently-mutated genes using `oncodriveFM`. Figure 3 shows the workflow that performs these tasks and produces various visualizations of the resulting gene set. Protein-centric lollipop plots, produced by a tool built on the `maftools` R package ^15^, facilitate visual recognition of patterns indicative of tumor suppressor genes and can also reveal mutation clustering and, in the extreme example, hot spots (e.g. *TMEM30A* and *NFKBIE*). Mutations in the latter two genes have previously been associated with DLBCL including a recurrent frameshift inducing indel observed here ^16,17^. *SPEN*, in contrast, has not been reported as recurrent target of somatic mutation in DLBCL but has been found mutated in other lymphoma types. The pattern of mutations suggests it may also act as a tumor suppressor gene in this cancer. We further noted *TET2*, *SETD1B*, *ARID1A*, *UBR5*, *DNMT3B* and *BTK* demonstrate similar mutation patterns. Although these have been identified as relevant genes in other cancers^18,19^, none of these have, to our knowledge, been previously ascribed to DLBCL.

**Figure 2.**
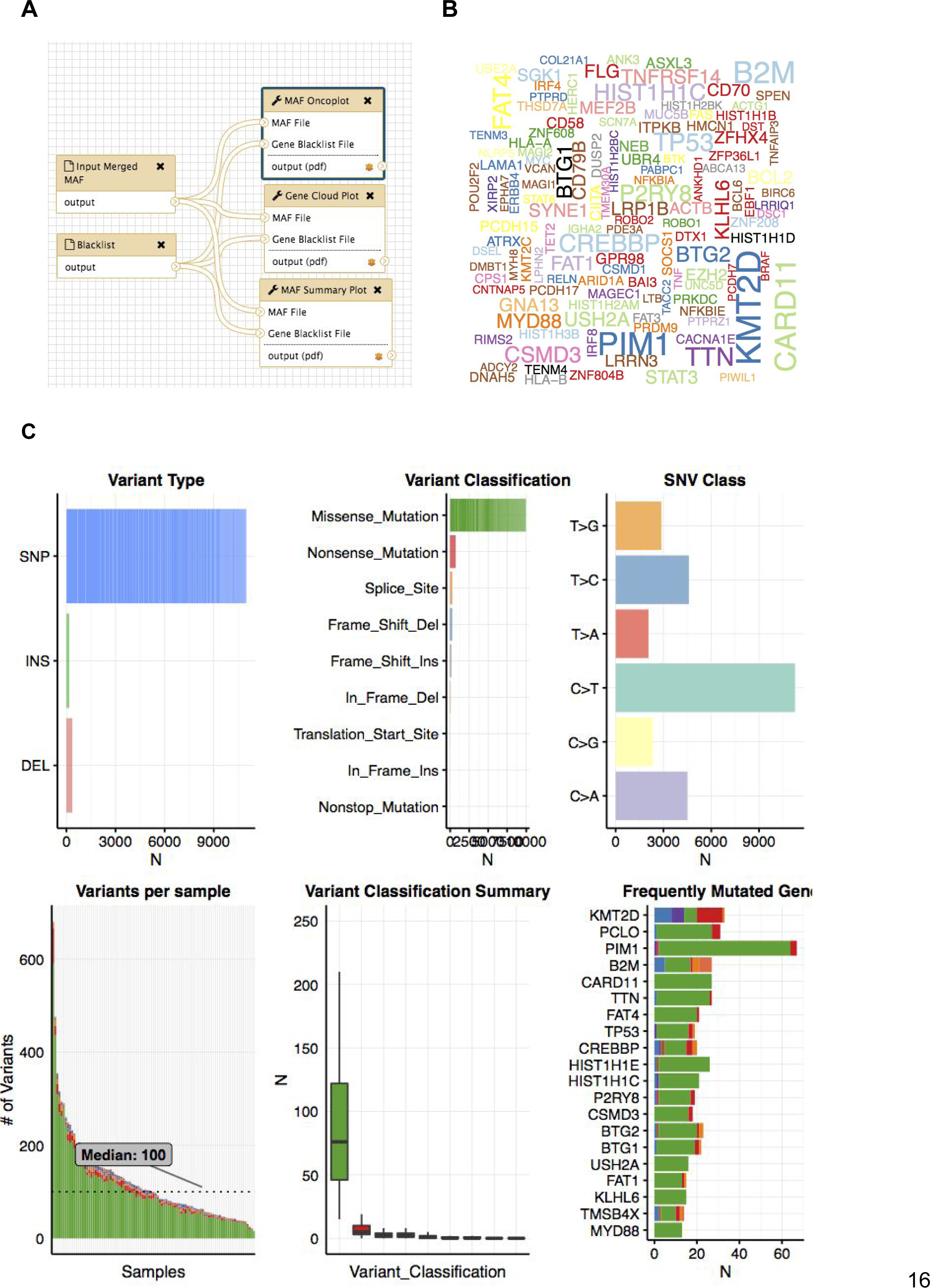
Cohort-wide summaries and visualization. Following primary mutation detection across a large cohort and annotation (i.e. with VEP using `vcf2maf`), it is useful to produce various summaries of the overall mutation burden and the types and classifications of mutations detected. The `maftools` R package offers a multitude of visualizations, many of which we have adapted into Galaxy. (A) In this example workflow, a merged MAF file containing the variants for the entire cohort of DLBCLs is input alongside a black-list of genes to hide from the outputs. (B) This word cloud, generated by the `genecloud` tool, provides a visually appealing summary of the frequency of mutations in genes above a user-specified threshold. (C) A generic `mafsummaryplot` tool provided by `maftools` generates six plots that represent descriptive features of the mutations and their annotations. It is evident that C>T is the predominant mutation type detected. A separate tool to perform refined mutation signature analysis is also available. Among the most commonly mutated genes are those previously attributed to DLBCL along with *TTN*, which encodes the largest human protein. With respect to the predicted effect, missense mutations are by far the dominant class of mutations. Despite this, tumor suppressors such as *KMT2D*, *TP53* and *B2M* show an elevation of inactivating mutation classes.

**Figure 3.**
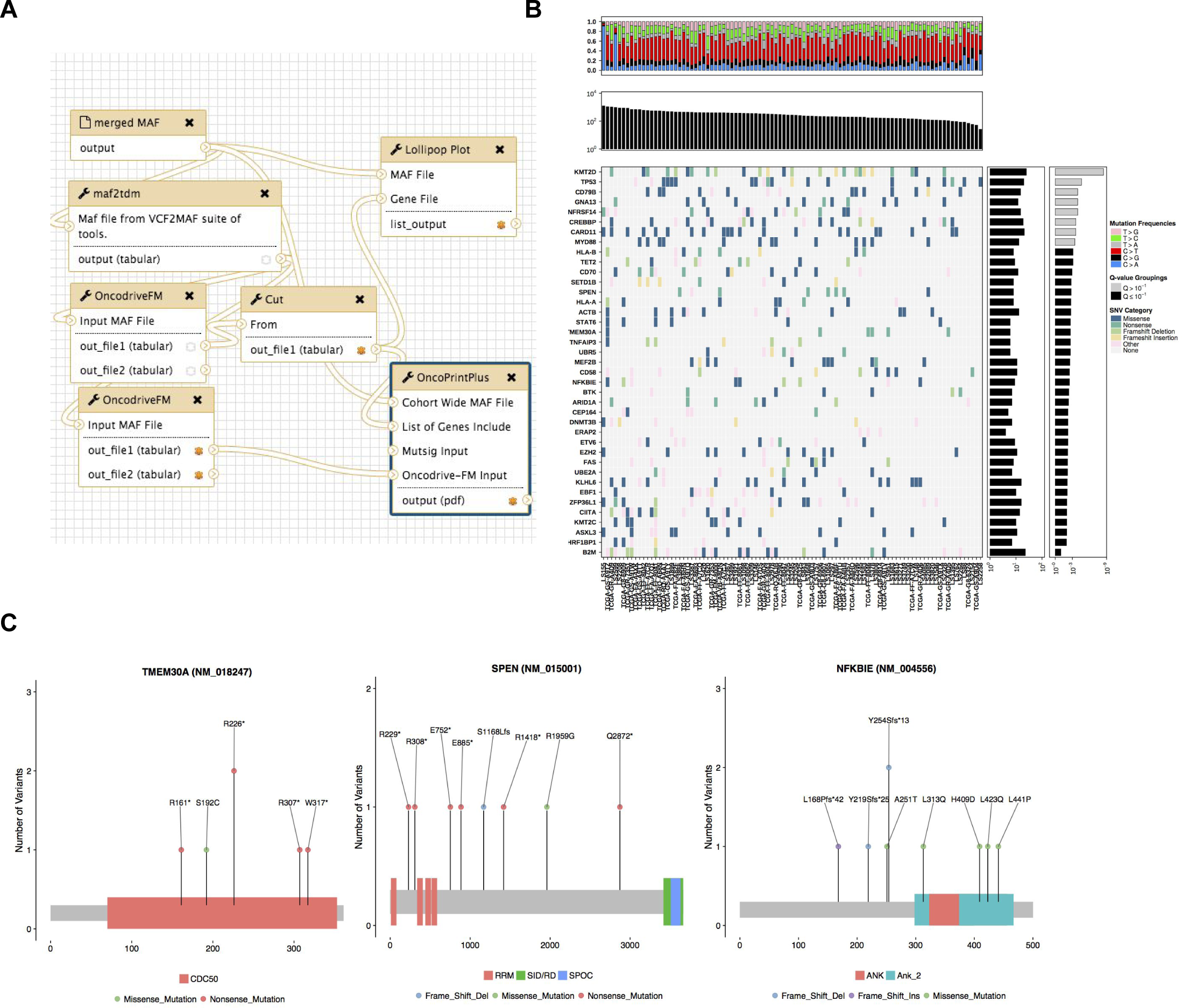
Significance analysis for mutation recurrence. (A) Tools have been implemented to screen mutation data for patterns of recurrence and identify significantly mutated genes. Shown above is an example workflow that utilizes the `OncodriveFM` algorithm and generates various visualizations for genes meeting a pre-specified Q-value threshold. (B) A common approach to summarize mutation data is a two-dimensional matrix with covariates plotted along the side axes. We implemented a tool that leverages multiplot in our R package to generate such images for arbitrary gene lists using the outputs of variant calling workflows that have been annotated using the `vcf2maf` tool. Mutations are colored based on the severity of mutations assigned automatically by the Ensembl Variant Effect Predictor (VEP) ^47^. Genes with more severe mutations are more likely to be tumor suppressor genes (e.g. *B2M* at the bottom and *TP53* and *KMT2D* at the top). Here, the total number of mutations detected in each patient is shown at the top and the P-value reported by `Oncodrive FM` is shown for each gene is shown on the right. The frequency of each of six possible mutation type can inform on mutational processes in individual samples. This is automatically determined from MAF files and is summarized at the top. (C) It is also often desirable to visualize the pattern of mutations within individual genes. The pattern is revealed using the `lollypopplot` tool that is run on each gene passing the threshold in this workflow.

We next attempted to integrate the exome-derived copy number information with the mutation calls. `Circos` is a popular approach to generate visualizations of genome-wide mutation data although it is generally better suited for genome-wide data and the representation of structural alterations and CNVs relative to genomic coordinates ^20^. For our application, we extended `Circos` to generate a gene-centric summarization of SNV and CNV data and produced the `Oncocirc os` Galaxy tool. Rather than plotting on a genomic coordinate scale, gene-level summaries of mutation data are mapped to their relative order on each chromosome and intergenic space (and genes with mutations below the threshold) are eliminated. To accomplish this, we implemented a custom parser that tabulates the data from MAF and segmented copy number files, applying a threshold to restrict the display to genes with a greater number of mutations cohort-wide. `Oncocircos` also accepts user-provided gene lists and regions of recurrent CNV (e.g. from `GISTIC`) and highlights these in the resulting image. Figure 4 shows the result of a workflow that runs `GISTIC` on a merged set of segmented data (in this example, from the `Sequenza` workflow) and integrates annotated SNV and indel calls from `Strelka`. In this visualization, several known DLBCL-associated recurrent events are observed including amplifications affecting *REL*, *MYC* and *BCL2* respectively on 2p, 8q and 18q. Recurrent deletions affecting the loci containing known tumor suppressor genes are also observable. A complementary visualization of these data is a gene by patient `oncostrip` in which annotated copy number and point mutations can be represented (Figure 6 and Supplemental Figure S4).

**Figure 4.**
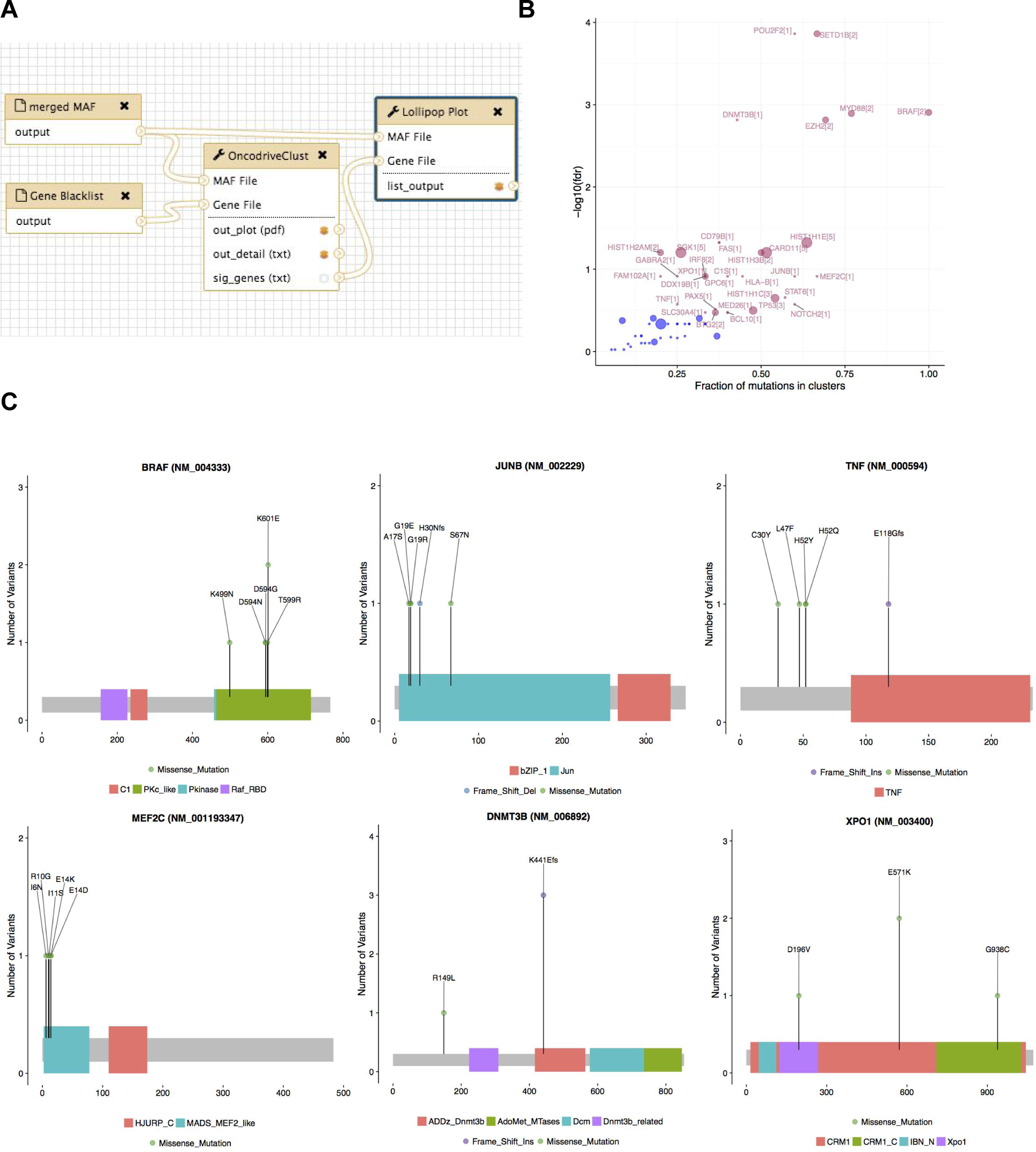
Identifying genes containing clustered mutations and hot spots. With sufficiently large cohorts, the pattern of non-silent mutations within the protein can inform on genes under specific selective pressure. A clear pattern seen in many dominantly acting cancer genes are mutation hot spots. The `OncodriveClust` workflow searches for genes with significant clustering of mutations that may represent hot spots or regions/sites whose mutation may produce a dominant effect. Application of this workflow (A) detected many lymphoma-related genes known to harbor mutation clusters (B). The workflow automatically generates lollipop plots for all genes above a user-specified FDR (in this example, 0.3) (C). Clear patterns of hot spots or mutation clusters are visible in each of these genes with only *BRAF* and *MEF2C* having been previously attributed to some DLBCLs ^21^.

## Discussion

### Towards reproducible and distributable workflows for cancer genome analysis

Large-scale efforts to understand the diversity of cancer-associated somatic alterations across common cancer types are continually expanding in scope. Many such efforts release raw (or aligned) tumor and normal sequence data into controlled-access repositories such as dbGAP and the European Genome-Phenome Archive (EGA). Owing to the many options and variations available in analytical methods, the mutations and copy number results presented along with these data are not directly amenable to direct comparisons between studies or pooled meta-analyses. Instead, the raw data must be obtained and processed uniformly alongside any new data sets. In light of the limited computational resources available to many research labs interested in incorporating existing sequence data into their analyses, some of the major data repositories are beginning to grant users permission to process patient data using cloud resources.

Availability and usability of analytical software are both critical factors in driving their adoption and the ultimate discovery of novel hallmarks. Accordingly, we provide a series of solutions that should accelerate adoption of our toolkit. First, providing automatic installation for tools wherever possible allow seamless integration into custom Galaxy instances. Second, many of the tools and workflows included here can be optionally configured to efficiently parallelize tasks on a cluster environment. Third, we show that our toolkit can be readily deployed onto a cloud-based Galaxy instance thereby eliminating the need for permanent access to commodity computing hardware. Together, this offers the potential to enable reproducible cancer research by empowering researchers to perform their own cancer genome analyses with unprecedented accessibility and directly share their workflows such that analyses can be reproduced on additional datasets by other groups.

Herein, we have used our toolkit to demonstrate the potential for knowledge discovery and hypothesis generation in a large-scale analysis of exome data. By summarizing cohort-level mutations with multiple complementary visualization tools, we have detected potential mutation patterns and novel candidate lymphoma-related genes. Such approaches can enable the discovery of signatures associated with specific molecular subgroups such as the two known molecular subgroups in DLBCL, namely activated B-cell (ABC) and germinal centre B-cell (GCB) varieties. Of note, each of the `oncoprintplus` and `oncostrip` tools now allow a variety of metadata, for example molecular subgroup, to be overlaid and these can be used to order samples (e.g. Figure 5).

**Figure 5.**
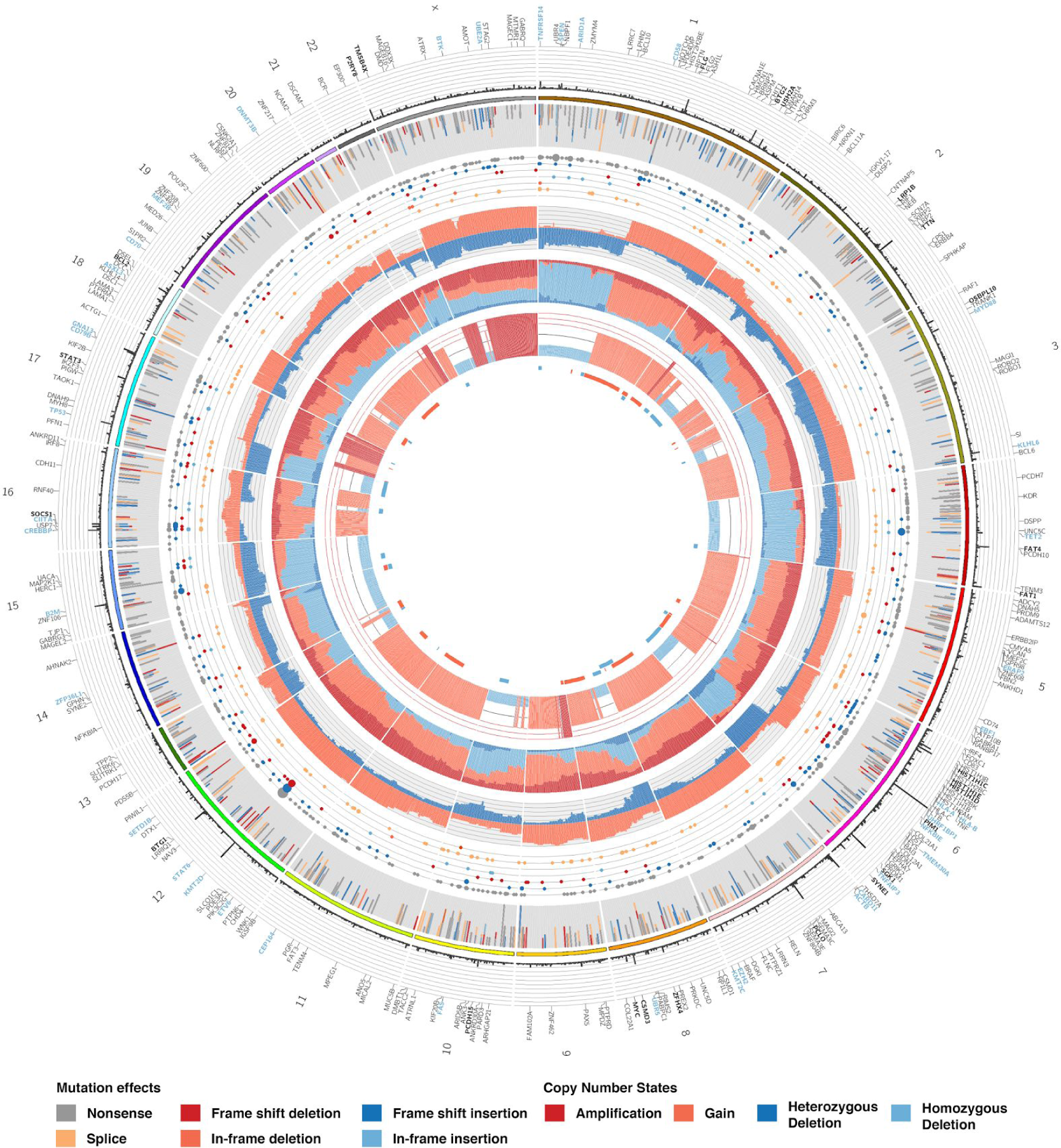
Visualization and data integration with Oncocircos. The new `Oncocircos` tool allows visualization of segment data derived from the Titan and Sequenza-based workflows we implemented. Genes exceeding a user-specified mutation frequency across the cohort are displayed and labels are automatically added for top genes. Those with at least twice the minimum mutation threshold are labeled in bold and those in an optional user-specified list can also be colored. A black-list file can be optionally provided to hide genes known to be enriched for artifacts. Stacked bar plots and circles provide summary of the annotated SNVs in each gene and a summary of the copy number state of each gene is provided in three inner tracks.

### Enabling new insights into DLBCL biology

The combined workflows employed here leverage distinct aspects of mutational information that can be individually leveraged to identify candidate cancer drivers and further integrated to inform on disease biology. Using a combination of methods, we provide additional evidence for the importance of several that have been attributed to DLBCL with weak support to date and those whose role as an oncogene or tumor suppressor has not been elucidated. For example, we note a hot spot in *NFKBIE* that induces a frameshift, which we recently observed in a separate set of patients with relapsed DLBCL ^17^. By employing `oncodriveclust`, we identified several genes with significant evidence for mutational recurrence. Mutations around the V600E hot spot in *BRAF* and within *MEF2C* have previously been reported to be present, albeit rare, in DLBCL ^21^. Another mutation we found to harbor a hot spot was STAT6 which, until recently, was thought to be mutated only in some less aggressive lymphomas such as FL and primary mediastinal B-cell lymphoma (PMBCL)<u>(Morin et al. 2016)</u>. A hot spot mutation in *XPO1* was also observed here. This mutation has recently been suggested as a molecular marker of PMBCL distinguishes it from true DLBCLs ^22^. One of the two cases bearing the canonical mutation (E571K) was among the few TCGA cases known to be PMBCLs and the other was from the second cohort for which clinical data was unavailable. These observations may further support the presence of mutations that will facilitate detection of PMBCL cases that can be difficult to distinguish from DLBCL.

The integration of mutation with copy number data using our tools has further informed on the potential relevance of some candidate lymphoma-related genes. *TMEM30A* demonstrated a mutation pattern indicative of tumor suppressor function (Figure 3C) and inspection of the `oncocircos` image suggest it resides within the commonly deleted region on 6q. Similarly, *FAT1* appears to have a strong signature towards inactivation and resides in a substantially smaller region that is commonly lost. Such patterns can be more readily confirmed using a separate visualization tool, namely `oncostrip` (Supplemental Figure S4). In contrast, some of the significantly amplified regions of the genome do not appear to harbor genes with significant evidence for recurrent mutations. Amplifications that include *JAK2* are known to be relevant to PMBCL but are not typically considered a feature of DLBCL. Upon inspection of the clinical data available for TCGA cases, we note that each of the four PMBCLs in this cohort contain a mutation or deletion affecting FAT1 and a JAK2 amplification. *POU2AF1*, which resides on 11q23.1, is a candidate target for the amplification of this region despite a low number of non-silent mutations and has been reported as commonly amplified in treatment-refractory DLBCLs ^23^. Further studies that include larger cohorts and possibly whole genome sequence data should help confirm the relevance of these observations.

Many of the genes known to be relevant to DLBCL biology are more commonly mutated in only one of the two molecular subgroups. Figure 6 shows the mutation distribution across some of these genes in the meta-cohort analyzed here, which has been organized on the predicted subgroup of each patient. Using the `oncostrip` tool to order patients on this designation uncovers additional genes in which mutations may be more common in the GCB subgroup such as *NFKBIE*, *ARID1A*, *FAS* and *STAT6*. *NFKBIE* mutations have recently been reported to be particularly common among PMBCLs and a marker of poor prognosis in that disease ^24^. One of the *NFKBIE* mutations detected herein was in a PMBCL case whereas the remainder were in nodal DLBCL cases and was almost exclusively seen in cases with other mutations suggestive of the GCB subgroup. This indicates a potential unappreciated role of *NFKBIE* in DLBCL or, taken together with our observation of mutations in *STAT6* and *XPO1*, may suggest that a significant subset of PMBCL cases may masquerade as GCB DLBCL. Further refinement of the mutation patterns of the two subgroups of DLBCL and PMBCL using larger cohorts is clearly warranted.

**Figure 6.**
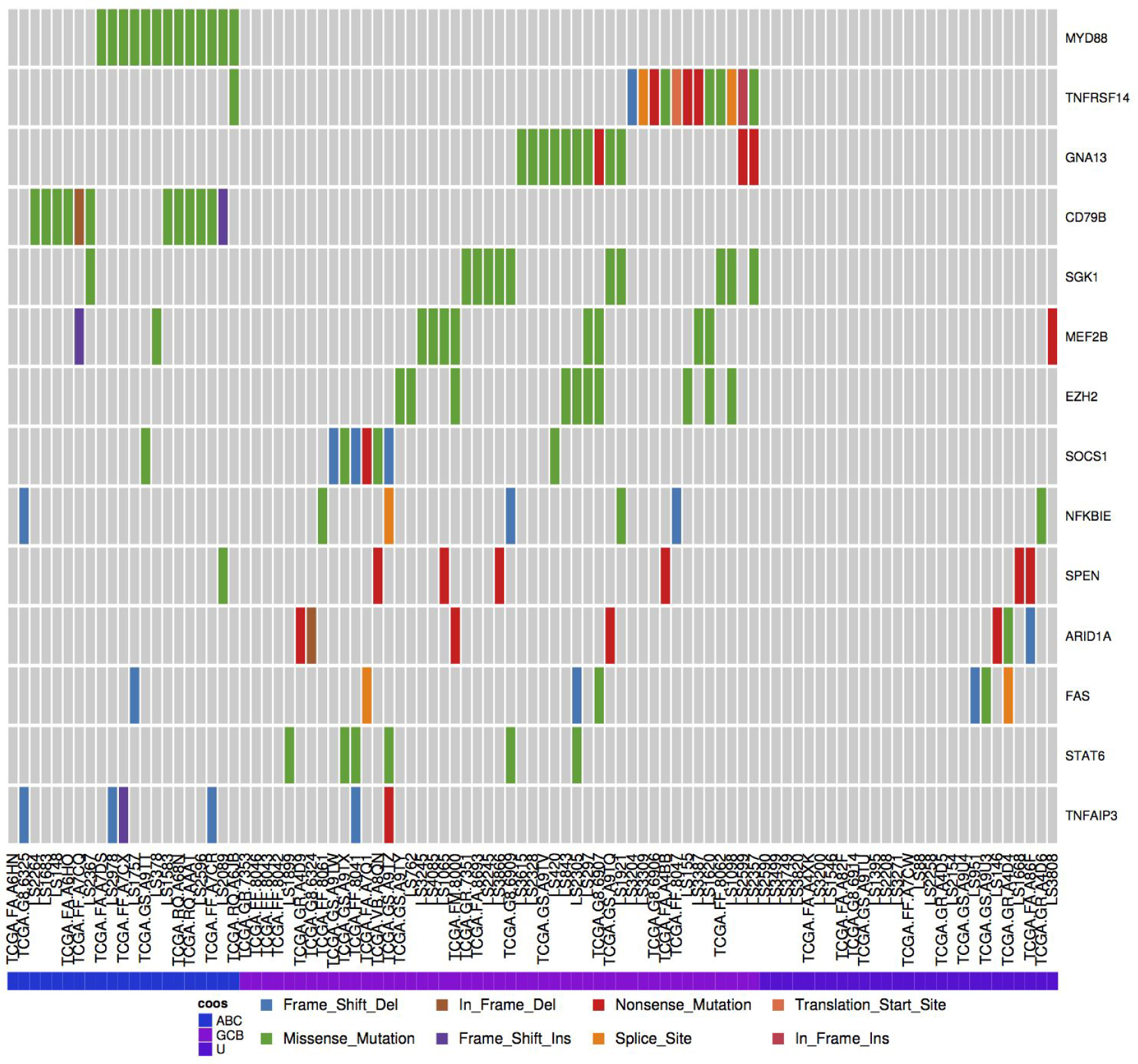
Discerning mutation patterns and identifying subtype-associated genes. DLBCL cases were assigned to either the ABC or GCB molecular subgroups using the presence of mutations known to be significantly restricted to either. Cases with no mutations unique to either molecular subgroup were designated unclassifiable (U).

## Conclusions

Our toolkit provides a growing list of standard methods for cancer genomic analysis and facilitates their deployment in a simplified, reproducible and accessible manner using Galaxy. We have successfully run our tools and workflows using AWS cloud computing and CloudMan, which provides an cluster environment to any research lab. We continue to provide new tools by extending functionality and updating versions as algorithms are refined. We also note that many of our tools have also been tested on whole genome sequence data and additional tools for performing analytical tasks better suited to that data type have been implemented but were not described in detail here. To facilitate scaling of these applications to such applications and to accelerate the analysis of exomes, we established new methods to accomplish parallelization in Galaxy. Overall, the release of this suite of tools provides the methods essential to drive discovery and eliminate the bottleneck in cancer genomic analysis.

## Methods

To integrate individual tools into Galaxy, we implemented XML-based configuration files which builds the command based on user-specified parameters. These contain a consistent design across tools and were developed using Planemo to ensure best practices ^25^. All repositories are stored on the main public Galaxy toolshed, which allows users to automatically install any tool ^26^. Modular tool dependency repositories provide the step-by-step instructions to compile necessary software for each tool. Previously defined repositories were recycled if available. We could not successfully produce tools that automatically install on all platforms but many of the tools successfully install (with dependencies) on the standard Galaxy AWS image and in a custom Ubuntu installation (v16.04). Synthetic alignment data containing artificial variants were generated and bundled with variant callers to enable automatic testing ^27^. To handle reference data, we have developed new Galaxy data managers but also allow the option of user-provided reference data ^28^.

We invested substantial effort to ensure that tools are parallelizable on cluster environments wherever it was deemed desirable and possible. Following the addition of new data types in the Galaxy codebase, this was subsequently re-implemented using the more transparent and efficient method that exploits the more recent Galaxy feature known as “data collections”. Briefly, parallelization of a workflow is accomplished by a combination of tasks (Figure 1), beginning with `fetch_interval`. This obtains chromosomal information from each input read alignment file and creates a collection of BED files defining all intervals available to each tool. To balance the load across all concurrently spawned jobs, we automatically pair large and small intervals. The second stage is a `preprocess` tool which defines all necessary preprocessing steps in a single tool and all will be executed together to reduce the numerous outputs associated with running multiple separate preprocessing tools in Galaxy. This includes a samtools flag and mapping quality filter, samtools remove duplicates and `bamutils clipoverlap`. The third stage involves running the selected tool on each of the intervals, allowing Galaxy to spawn processes to available CPUs. The fourth stage is `postprocess`, which follows similar methodology to `preprocess`. Example usage includes further variant filtration and annotation steps. Finally all output files are merged, if necessary, so they may be supplied to subsequent tools and workflows. For tools that can be multithreaded, we instead leverage this capability instead of using chromosomal splitting.

Many of the analyses and workflows shown in detail in Figures 1-6 were performed on a local Galaxy instance on a linux-based server. For computationally demanding tasks, we launched a Galaxy instance on AWS Elastic Cloud Compute (EC2) using CloudMan and installed the workflow and tool dependencies. A cluster configuration consisting of one r3.8xlarge master node and five r3.2xlarge worker nodes was selected. We successfully uploaded 96 bam files representing the cohort of published DLBCL samples and ran all SNV and CNV workflows on each tumour/normal pair and captured details on runtime and speedup associated with parallelization (not shown) ^21^.

## List of Abbreviations

SNV: :Single Nucleotide Variant
SV: :Structural Variant
CNV: :Copy-Number Variant
GUI: :Graphical User Interface
AWS: :Amazon Web Services

## Competing Interests

The authors declare they have no competing interests.

## Authors’ Contributions

M.A.A. and E.R. were responsible for deploying tools in galaxy. M.A.A. and B.M.G. benchmarked workflows on AWS. M.A.A. and M.K. created figures. M.A.A., B.M.G. and R.D.M. wrote the manuscript, which was reviewed and approved by all authors. R.D.M, P.C.B. and S.P.S. led the study.

## Acknowledgements

The results published here are in whole or part based upon data generated by the TCGA Research Network: http://cancergenome.nih.gov/. We gratefully acknowledge TCGA and all providers of samples and resources for generating this valuable resource. The TCGA exome data was obtained through dbGAP (phs000178.v9.p8 and phs000450.v2.p1) and the latter has been described previously ^21^. Said data was produced as part of the Slim Initiative for Genomic Medicine (SIGMA), a joint U.S.-Mexico project funded by the Carlos Slim Health Institute. This work was supported by a contract from Genome Canada and Genome British Columbia (173CIC). funding from Mitacs (awarded to Morin) and Amazon AWS research grant. Sequencing of the large DLBCL cohort was funded by an operating grant from CIHR (to RDM). RDM is supported by New Investigator Awards from the Canadian Institutes for Health Research and the Terry Fox Research Institute. We thank all members of the Boutros, Shah and Morin research groups for feedback on this work. We also thank the Galaxy community for their ongoing support. We are particularly grateful to Enis Afgan, John Chilton, Nitesh Turaga and Björn Gruening for their gracious assistance. We also thank Marija Jovanovic for assisting in tool deployment.

---

**Supplementary Table S1.** Helper tools implemented to facilitate tool linkage and parallelization.

| Tool name | Description | Application |
| --- | --- | --- |
| ensemble_vcf | An ensemble variant caller that applies a voting scheme | Select high-confidence SNV calls from the outputs of multiple variant callers |
| augment_maf | Add allele counts for each variant from tumour and normal bam files and integrate mutations from multiple samples. | A key step in the process in preparing input for PyClone |
| ensembl_vep | Annotate effect of SNVs and indels using the Ensembl build | Used by vcf2maf to produce annotated MAF files from raw mutation calls in VCF format |
| fetch_interval | Reads a Binary Sequence Alignment File (BAM) and writes out intervals in tab-delimited format | Preprocessing in parallelization |
| merge, merge_gzip | Merge while maintaining order of text files in a data collection | Postprocessing in parallelization |
| merge_maf_collection | Merge a collection of MAF files and handle headers | Creating cohort-wide MAF file from individual samples |
| preprocess |  |  |
| cnv2igv | Reformat outputs from CNV callers into standard IGV-friendly input | Produce inputs for GISTIC and Oncocircos tools |
| igv2gistic | Incorporate exon annotations to produce segmented data and marker file | Preprocessing necessary to run GISTIC2.0 on exome-derived CNV data |
| select_optimal_cluster | Nominate Titan run from a sample with the best fit | Postprocess Titan output to yield best result for further analysis |

**Supplementary Figure S1.**
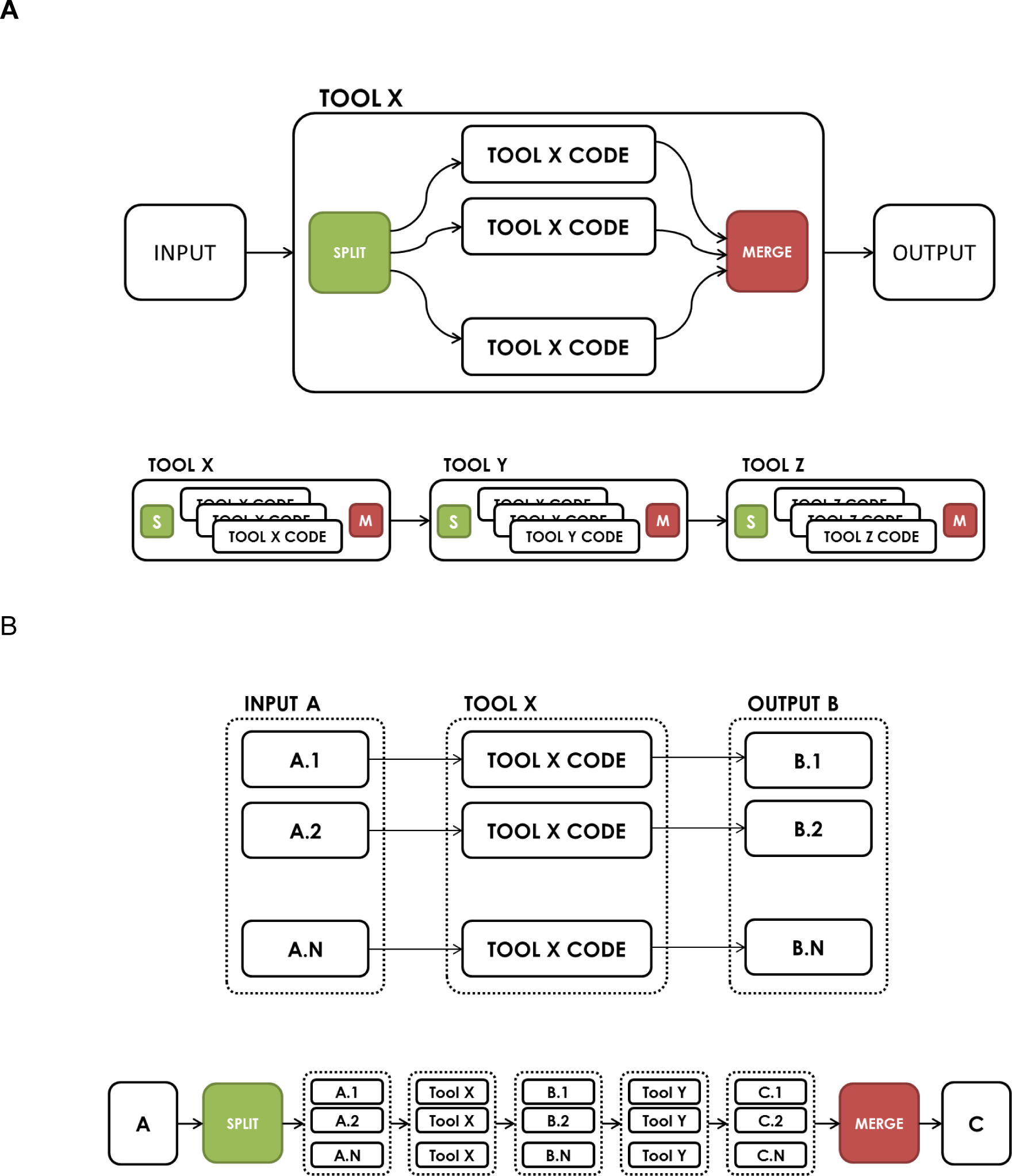
Achieving parallelization in Galaxy. There are two ways to achieve parallelization in Galaxy. The first (A) employs the parallelism tag, which calls specific split and merge functions depending on the galaxy input and output data type. These functions are predefined within galaxies codebase. The second (B) uses galaxy collections, which are essentially containers of input files. Inputs can be split into a collection of files and subsequently pipeline these through a series of tools. When complete, the individual outputs can be merged. Parallelizing using collections is far more transparent to the user and also limits that number of unnecessary split and merge functions.

**Supplementary Figure S2.**
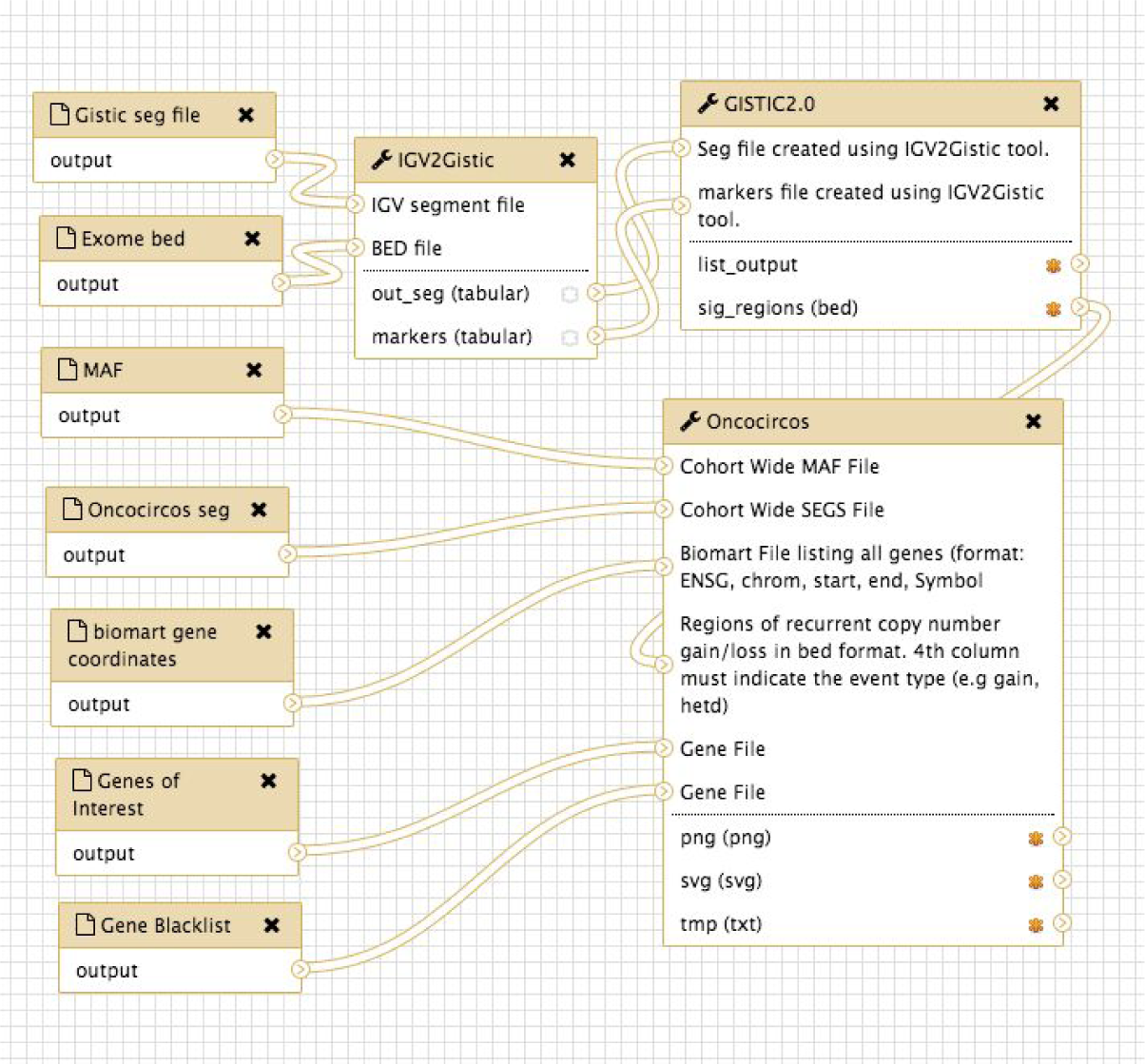
An ensemble approach to detect somatic SNVs. The `ensembl_vcf` tool receives the output of variant callers and selects variants detected by a user-specified number of tools. This example workflow runs four variant callers (`strelka, mutationSeq, RADIA` and `SomaticSniper`) and runs `vcf2maf` to annotate the resulting list of variants with support from a sufficient number of tools.

**Supplementary Figure S3.**
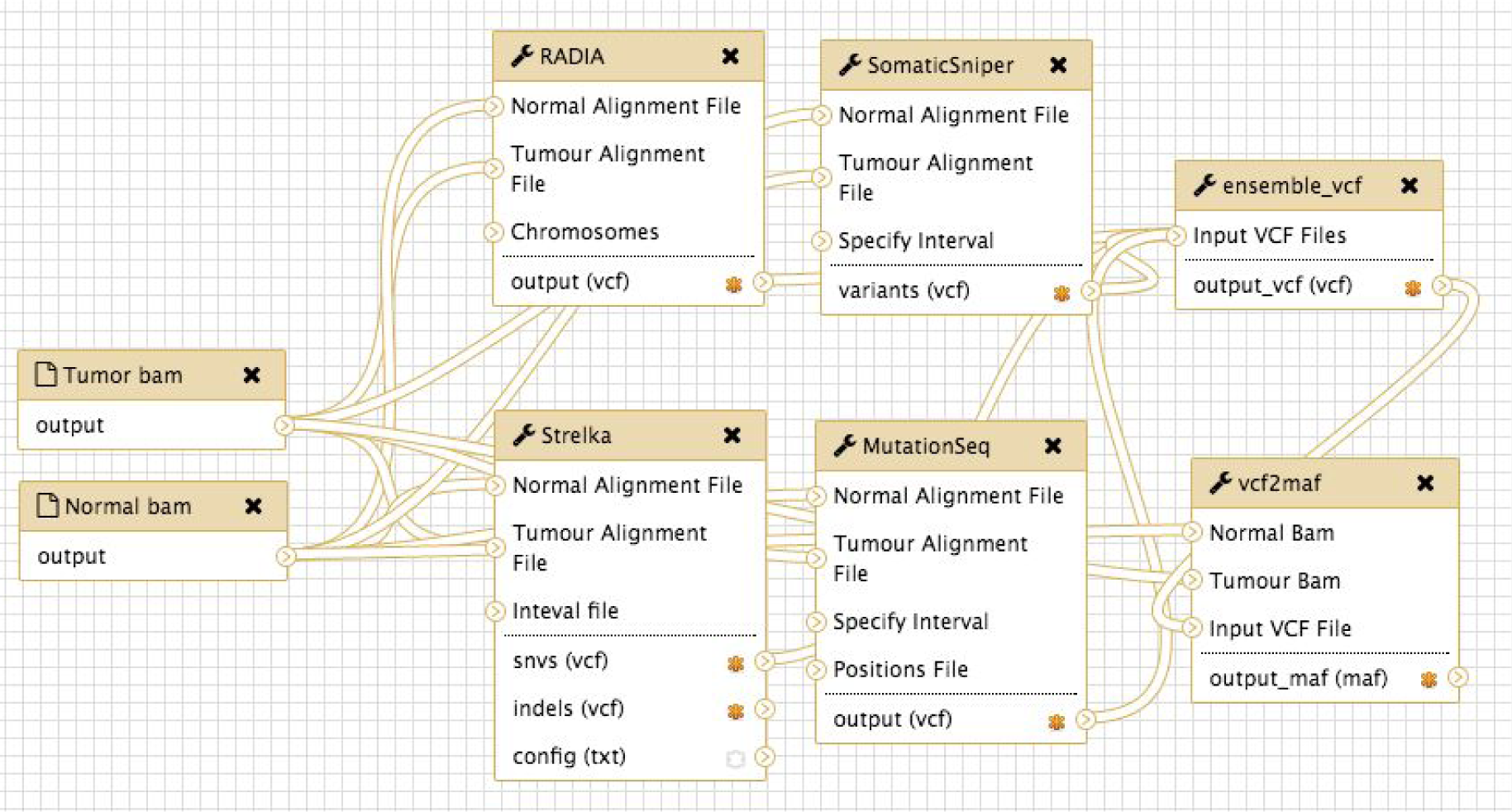
A workflow to integrate SNV and CNV data and produce integrative visualizations. This workflow uses exome-derived CNV and SNV data to generate a list of recurrently gained/lost genomic regions (using `GISTIC`) and displays these along with gene-centric summaries of segmented copy number and SNV data using `Oncocircos`. To generate Figure 5, we included a blacklist containing all immunoglobulin genes and Mucin genes whereas the genes identified as significantly mutated by `oncodriveFM` were provided separately to enforce highlighting.

**Supplementary Figure S4.**
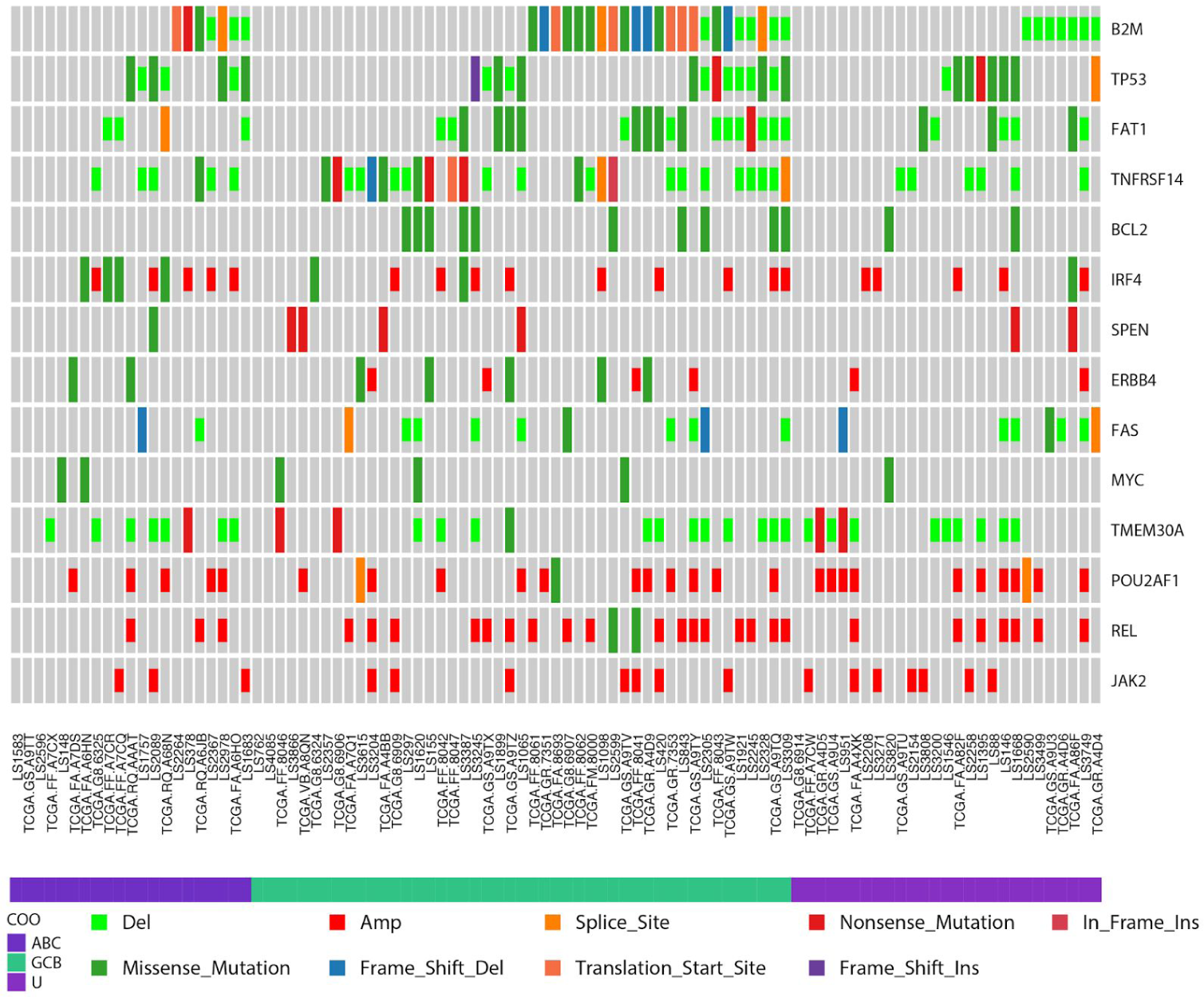
Using the Oncostrip tool to integrate copy number and mutation data. It can be desirable to visualize the complete set of mutational information cohort-wide without losing the patient-mutation relationships and potential gene-gene interactions that are not retained in `Oncocircos`. For this application, the `oncostrip` component of `maftools` can also accept raw outputs from `GISTIC`. Here, we have included the known gene targets of some recurrent amplifications and deletions detected in the cohort (*REL*, B2M and *TNFRSF14*). Each of *FAT1* and *TMEM30A* reside in significantly deleted regions and bear a combined pattern of mutation and deletion consistent with tumor suppressor function.

